# Functional validation of *EIF2AK4* (GCN2) missense variants associated with pulmonary arterial hypertension

**DOI:** 10.1101/2024.01.27.577559

**Authors:** Giulia Emanuelli, JiaYi Zhu, Nicolas W Morell, Stefan J Marciniak

**Author notes:** Correspondence to: Stefan J. Marciniak, Cambridge Institute for Medical Research (CIMR), University of Cambridge School of Clinical Medicine, The Keith Peters Building, Cambridge Biomedical Campus, Hills Rd. Cambridge, CB2 0XY.

## Abstract

Pulmonary arterial hypertension (PAH) is a disorder with a large genetic component. Biallelic mutations of *EIF2AK4*, which encodes the kinase GCN2, are causal in two ultra-rare subtypes of PAH, pulmonary veno-occlusive disease and pulmonary capillary haemangiomatosis. *EIF2AK4* variants of unknown significance have also been identified in patients with classical PAH, though their relationship to disease remains unclear. To provide patients with diagnostic information and enable family testing, the functional consequences of such rare variants must be determined, but existing computational methods are imperfect. We applied a suite of bioinformatic and experimental approaches to sixteen *EIF2AK4* variants that had been identified in patients. By experimentally testing the functional integrity of the integrated stress response (ISR) downstream of GCN2, we determined that existing computational tools have insufficient sensitivity to reliably predict impaired kinase function. We determined experimentally that several *EIF2AK4* variants identified in patients with classical PAH had preserved function and are therefore likely to be non-pathogenic. The dysfunctional variants of GCN2 that we identified could be subclassified into three groups: misfolded, kinase-dead, and hypomorphic. Intriguingly, members of the hypomorphic group were amenable to paradoxical activation by a type-1.5 GCN2 kinase inhibitor. This experiment approach may aid in the clinical stratification of *EIF2AK4* variants and potentially identify hypomorophic alleles receptive to pharmacological activation.

## Introduction

Aberrant vascular remodelling in pulmonary arterial hypertension (PAH) raises pressures in the pulmonary vasculature to cause right heart failure^1^. Affected young adults often suffer progressive disease leading to premature death. Although classical PAH is most frequently caused by mutations in the TGFβ/BMP signalling axis^2–5^, rare subtypes such as pulmonary veno-occlusive disease (PVOD) and pulmonary capillary haemangiomatosis (PCH) have distinct genetics and are less amenable to current clinical management^6^. With no effective treatments apart from lung transplantation, death occurs within a year in 72% of patients diagnosed with these aggressive PAH subtypes^7^.

Since the first report in 2014 linking biallelic mutations of *EIF2AK4* to PVOD^8^, approximately one hundred *EIF2AK4* alleles have been reported to be associated with PAH and its subtypes^4,8–14^. Although frameshift mutations constitute a large proportion, approximately a third (34) of these alleles are missense variants, the functional consequences of which are unknown (Figure 1)^15^. Validating the pathogenicity of such variants of uncertain significance (VUSs) would aid in diagnosis, enable cascade genetic testing of relatives, and recruitment of patients to chemoprotective clinical trials^6^.

**Figure 1.**
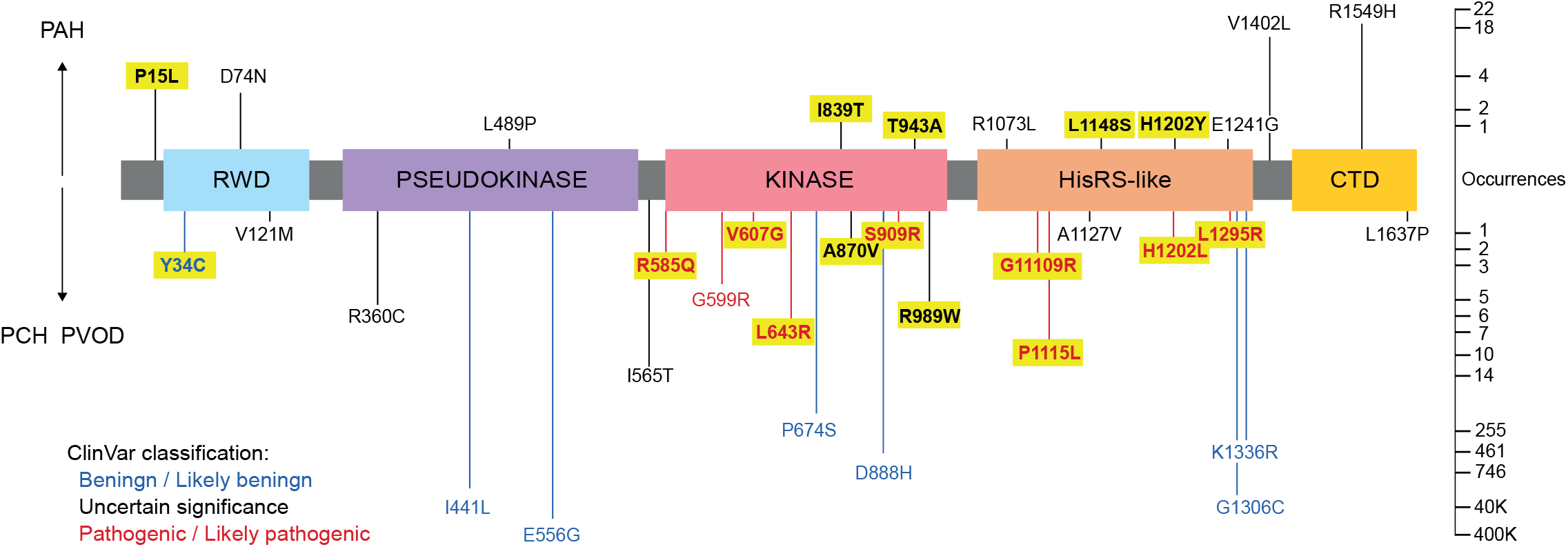
Schematic of known patient-specific missense variants. Variants over the domain schematic were from patients with classical PAH. Variants below were from patients with PCH and PVOD. Variants reported to be benign/likely benign in ClinVar database^*33*^ are in blue, those reported as pathogenic/likely pathogenic are in red, black represents variants of uncertain significance. Occurrences represent cumulative allele counts in the gnomAD database^*34*^ or published reports^8–14^. Highlighted variants are analysed experimentally in this study.

*EIF2AK4* encodes GCN2, a large serine/threonine kinase homodimer that responds to amino acid depletion by monitoring the efficiency of protein synthesis through its interaction with stalled ribosomes^16,17^. GCN2 is comprised of an N-terminal RWD domain (120-137) involved in protein-protein interactions, pseudokinase (276-539) and kinase domains (585-1016) including GCN2’s dimerisation interface, an HisRS-like domain (1058-1490) and C-terminal domain (1533-1649) where binding to ribosomes and recognition of uncharged tRNAs occur^18^. When activated, GCN2 triggers a cellular signalling pathway termed integrated stress response (ISR) by phosphorylating the α subunit of eukaryotic initiation factor 2 (eIF2α)^19,20^. This attenuates most protein synthesis, while enhancing the translation of ISR-specific mRNAs, such as that encoding the transcription factor ATF4, owing to the presence of upstream open reading frames (uORFs) in their 5’UTRs^21^. An eIF2α phosphatase subunit, PPP1R15A, is similarly regulated and its expression eventually terminates the ISR^20–22^. Due to its fundamental biological roles, disruption of the ISR is implicated in many diseases^23–25^.

Technological advances have made genomic sequencing readily available in the clinic, leading to a proliferation in the number of VUSs encountered. Currently, strategies to predict the impact of novel genetic variants are largely restricted to computational methods that rely on evolutionary conservation (e.g. SIFT, PolyPhen)^26^ or integrate a range of scores from existing predictive tools (e.g. CADD, REVEL)^27,28^. Other methodologies include the computation of folding free energy differences (FoldX)^29^, which predicts protein stability. More recently, deep learning approaches have been developed (EVE, AlphaMissense)^30,31^ that account for both evolutionary constraints and structural information. Although these have improved predictive accuracy, computational approaches remain imperfect. We set out to examine patient-specific *EIF2AK4* missense mutations both *in silico* and experimentally using existing bioinformatic tools and cell biological assays. In so doing, we subclassified PAH-associated GCN2 variants in functional (likely benign), destabilised/misfolded, or kinase impaired. A subset of the kinase impaired variants showed preserved target engagement. These hypomorphic variants were amenable to paradoxical activation using an ATP-competitive GCN2 inhibitor^32^. Based on these results, we propose a simple methodology for the experimental validation of the functionality of *EIF2AK4* VUSs, which outperforms existing computational approaches.

## Results

### Computational analysis of PAH-associated *EIF2AK4* missense variants suggests heterogeneity

Thirty-four missense variants of *EIF2AK4* (RefSeq: NC_000015.10) have been published^8–14^ or reported in ClinVar^33^ to be associated with PAH (Table 1). Genome sequencing of the general population^34^ suggests that most of these variants are rare with allele frequencies < 0.01, though three, I441L, E556G and G1306C, are common with frequencies > 0.1 (Figure 1). Nine have already been classified as pathogenic/likely pathogenic (R585Q, G599R, V607G, L643R, S909R, G11109R, P1115L, H1202L and L1295R) and seven as benign/likely benign, including the three common variants (Y34C, I441L, E556G, P674S, D888H, G1306C and K1336R), but most remain uncharacterised VUSs (Figure 1, Table 1).

**Table 1.** Known pulmonary hypertension-associated missense variants of GCN2. ClinVar classification (VUS: variant of uncertain significance) and patient diagnosis where available: Hom = homozygous, Het = heterozygous, C Het = compound heterozygous. Scores SIFT (T: tolerated; NT: not tolerated), PolyPhen2, CADD v1.6, REVEL and AlphaMissense (high scores corresponding to more severe consequences). FoldX use to estimate difference in Gibbs free energy (ΔΔG, kcal/mol) between a variant and the non-mutated GCN2 (scores >1 indicate predicted destabilisation, scores >3 indicate predicted severe misfolding).

| DNA Base | Amino Acid | Pubication | ClinVar classification | Patient Diagnosis | SIFT | PolyPhen2 | CADDv1.6 | REVEL | Alpha Missense | FoldX: $\Delta\Delta G$ (kcal/mol) | Integrative *in silico* functional prediction |
| --- | --- | --- | --- | --- | --- | --- | --- | --- | --- | --- | --- |
| 44C>T | P15L | Hadinnapola 2017 | VUS | PAH (Het) | T | 0 | 22.6 | 0.133 | 0.0834 | 0.23 | Non-pathogenic |
| 145-2A>G | Y34C | UK cohort (unpublished) | Benign/likely-benign | PVOD (C Het) | T | 0.999 | 32 | 0.634 | 0.4589 | 3.79 | Misfolded |
| 220G>A | D74N | Hadinnapola 2017 | VUS | PAH (Het) | NT | 0.954 | 28 | 0.166 | 0.1726 | 0.38 | Non-pathogenic |
| 361G>A | V121M | Montani 2017 | VUS | PVOD (C Het) | T | 1 | 33 | 0.467 | 0.7125 | -1.04 | Non-pathogenic |
| 1078C>T | R360C | ClinVar RCV001287133 | VUS | PCH | NT | 0.999 | 26.3 | 0.566 | 0.083 | 1.68 | Uncertain |
| 1321A>C | I441L | ClinVar RCV000999785 | Benign/likely-benign | PCH | T | 0.001 | 16.9 | 0.075 | 0.0839 | -0.70 | Non-pathogenic |
| 1466C>T | L489P | Gómez 2015 | VUS | PAH | NT | 1 | 29 | 0.895 | 0.9737 | 7.89 | Misfolded |
| 1667A>G | E556G | ClinVar RCV000755517 | Benign/likely-benign | PCH | T | 0 | 22.5 | 0.088 | 0.0523 | 0.63 | Non-pathogenic |
| 1694T>C | I565T | ClinVar RCV001285279 | VUS | PCH | T | 0.835 | 22.2 | 0.307 | 0.2246 | -0.36 | Non-pathogenic |
| 1754G>A | R585Q | Eyries 2014; Montani 2017 | Pathogenic/likely-pathogenic | PVOD; PCH | NT | 1 | 32 | 0.861 | 0.9272 | 2.12 | Kinase-deficient |
| 1795G>C | G599R | Hadinnapola 2017 | Pathogenic/likely-pathogenic | PAH; PCH (Hom) | NT | 1 | 32 | 0.952 | 0.9991 | 0.83 | Kinase-deficient |
| 1820T>G | V607G | Hadinnapola 2017 | Pathogenic/likely-pathogenic | PAH (C Het); PCH | NT | 0.998 | 33 | 0.557 | 0.8039 | 4.73 | Misfolded |
| 1928T>G | L643R | Eyries 2014; Levy 2016; Montani 2017 | Pathogenic/likely-pathogenic | PVOD; PCH | NT | 0.999 | 32 | 0.889 | 0.9911 | 9.65 | Misfolded |
| 2020C>T | P674S | ClinVar RCV001286399 | Benign/likely-benign | PCH | T | 0.029 | 12.1 | 0.077 | 0.0694 | -1.45 | Non-pathogenic |
| 2516T>C | I839T | Hadinnapola 2017 | VUS | PAH (Het) | NT | 0.99 | 29.6 | 0.781 | 0.9854 | 2.90 | Non-pathogenic |
| 2609C>T | A870V | Montani 2017 | VUS | PVOD (C Het) | T | 1 | 28.1 | 0.658 | 0.9943 | 0.95 | Uncertain |
| 2662G>C | D888H | ClinVar RCV001287878 | Benign/likely-benign | PCH | NT | 0.998 | 24.8 | 0.237 | 0.1005 | 0.59 | Non-pathogenic |
| 2727C>G | S909R | Hadinnapola 2017 | Pathogenic/likely-pathogenic | PAH (Het); PCH | NT | 1 | 28 | 0.446 | 0.9985 | 5.67 | Misfolded |
| 2827A>G | T943A | Hadinnapola 2017 | VUS | PAH (C Het) | NT | 0.997 | 25.4 | 0.464 | 0.8597 | 1.54 | Non-pathogenic |
| 2965C>T | R989W | Eichstaedt 2022 | VUS | PVOD | NT | 1 | 28.9 | 0.862 | 0.9838 | 38.59 | Misfolded |
| 3218G>T | R1073L | Hadinnapola 2017 | VUS | PAH | NT | 0.995 | 32 | 0.466 | 0.3374 | 0.15 | Non-pathogenic |
| 3325G>A | G1109R | Hadinnapola 2017 | Pathogenic/likely-pathogenic | PAH (C Het); PCH | NT | 1 | 32 | 0.766 | 0.9856 | 9.20 | Uncertain |
| 3344C>T | P1115L | Gómez 2015; Tenorio 2015 | Pathogenic/likely-pathogenic | PAH; PVOD; IPAH; PCH | NT | 1 | 29.7 | 0.702 | 0.8415 | 1.46 | Uncertain |
| 3380C>T | A1127V | Eichstaedt 2022 | VUS | PVOD | T | 1 | 32 | 0.262 | 0.6049 | 6.08 | Misfolded |
| 3443T>C | L1148S | Eichstaedt 2022 | VUS | CTD-APAH | T | 1 | 28.4 | 0.341 | 0.2648 | 1.11 | Non-pathogenic |
| 3604C>T | H1202Y | Hadinnapola 2017 | VUS | PAH (Het) | T | 0.996 | 26.5 | 0.716 | 0.812 | -0.44 | Non-pathogenic |
| 3605A>T | H1202L | Hadinnapola 2017 | Pathogenic/likely-pathogenic | PAH; PCH (Hom) | NT | 0.998 | 28.4 | 0.802 | 0.9507 | 3.00 | Uncertain |
| 3722A>G | E1241G | Hadinnapola 2017 | VUS | PAH | NT | 0.971 | 33 | 0.342 | 0.7069 | 1.50 | Non-pathogenic |
| 3884T>G | L1295R | Hadinnapola 2017 | Pathogenic/likely-pathogenic | PAH; PCH (C Het) | NT | 1 | 28.8 | 0.814 | 0.9408 | 4.89 | Misfolded |
| 3916G>T | G1306C | ClinVar RCV000999798 | Benign/likely-benign | PCH | NT | 1 | 31 | 0.307 | 0.2596 | 2.26 | Uncertain |
| 4007A>G | K1336R | ClinVar RCV003103836 | Benign/likely-benign | PCH | T | 0 | 15.8 | 0.063 | 0.0564 | 0.14 | Non-pathogenic |
| 4204G>T | V1402L | Montani 2017 | VUS | IPAH | T | 0.997 | 24.7 | 0.285 | 0.4847 | -0.19 | Non-pathogenic |
| 4646G>A | R1549H | Hadinnapola 2017 | VUS | PAH (Het) | NT | 0.998 | 32 | 0.318 | 0.6517 | 0.30 | Non-pathogenic |
| 4910T>C | L1637P | Montani 2017 | VUS | PVOD (C Het) | NT | 1 | 31 | 0.624 | 0.9982 | 5.98 | Misfolded |
|  | K619R |  | - | kinase-dead | NT | 1 | 32 | 0.783 | 0.9769 | 0.20 | Kinase-deficient |

We applied existing computational methods including SIFT, PolyPhen2, CADDv1.6, REVEL and AlphaMissense in an effort to predict the functional significance of these GCN2 variants^26–28,30^ (Table 1). Higher scores represent higher predicted severity. We also used FoldX5.0 to estimate differences in Gibbs free energy (ΔΔG) between wildtype GCN2 and each variant (Table 1)^29^. Although broadly concordat, the algorithms yielded several discordant results. For example, although H1202Y was predicted to maintain stable folding by FoldX, and tolerated by SIFT and CADD, it was categorised as severe/likely pathogenic by the other methods. Conversely, Y34C was predicted to be destabilised by FoldX (ΔΔG>1.5kcal/mol) and likely-pathogenic by PolyPhen, CADD and REVEL, but tolerated by SIFT and categorised as ambiguous by AlphaMissense. Nevertheless, by integrating these computational methods we predicted missense variants of GCN2 to be either (i) benign, (ii) misfolded, or (iii) kinase deficient (Table1).

### Expression of GCN2 variants and ISR reporter activation

To test the *in silico* predictions, we next performed experimental validation in cultured cells. A bioluminescent ISR reporter was generated by fusing the 5’UTR of ATF4 with NanoLuc® luciferase (ATF4::NanoLuc, Figure 2A). We obtained optimal results when using human rather than murine ATF4, driven by a CMV promoter/SV40 enhancer (data not shown). uORFs in the 5’UTR of ATF4 mRNA impose translational regulation on the downstream coding sequence^21^. When endogenous GCN2 was deleted in HeLa cells (GCN2 KO), as expected activation of the ISR reporter by the histidyl-tRNA synthetase inhibitor histidinol was ablated (Figure 2B). Re-expression of wild type human GCN2, but not a kinase-dead mutant (K619R) rescued ATF4::NanoLuc responsiveness to histidinol, validating the system (Figure 2B).

**Figure 2.**
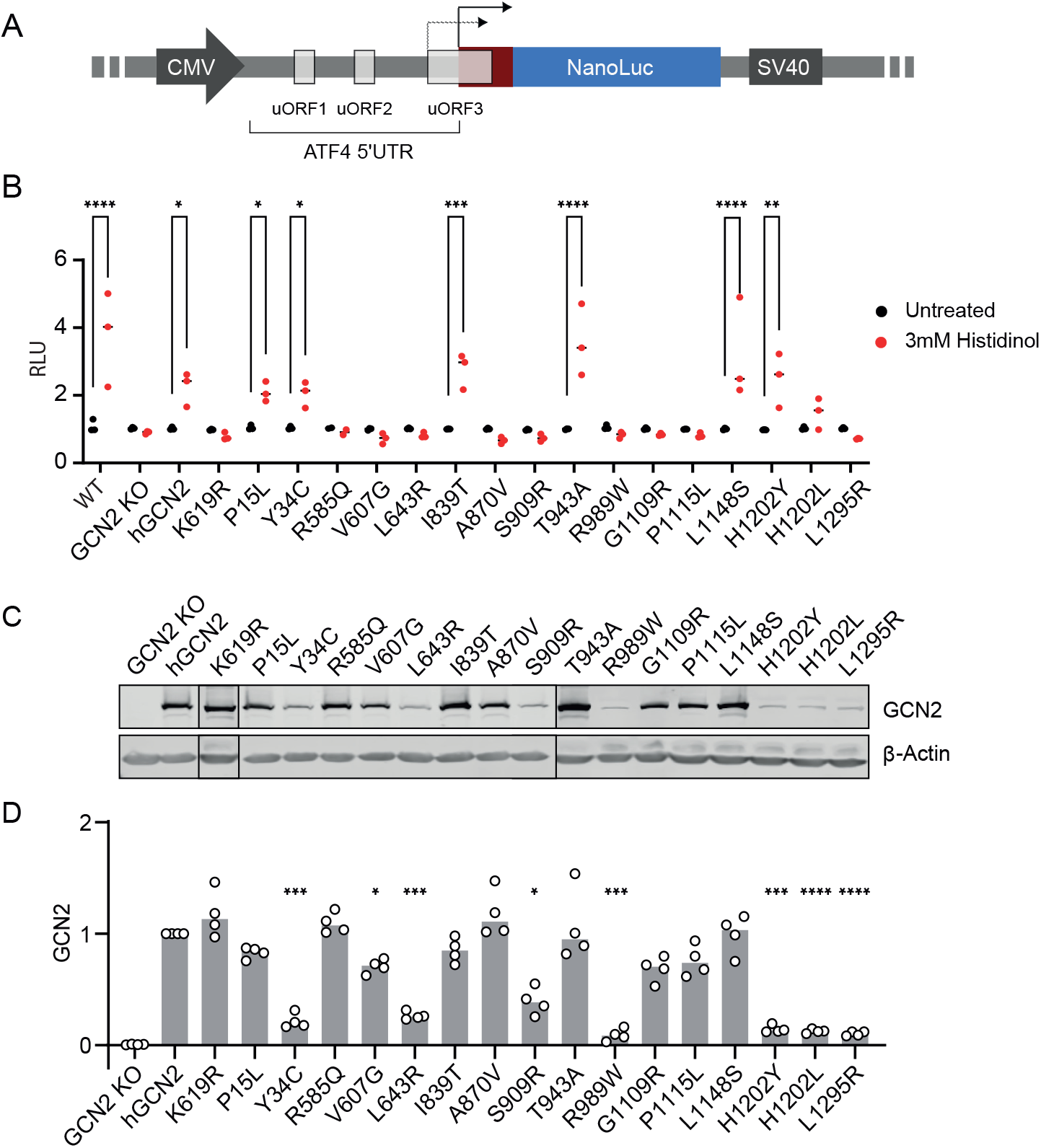
Variant in cell activity and expression. (A) ISR reporter construct comprising 5’ UTR of human ATF4 containing upstream open reading frames (uORF) cloned in frame with NanoLuc® luciferase. CMV promoter and SV40 enhancer. Under conditions of low eIF2α phosphorylation, uORF3 is translated resulting in low reporter signal. When eIF2α phosphorylation increases, ATF4 initiator AUG (maroon) is translated leading to NanoLuc reporter synthesis. (B) Reporter activity (relative light units, RLU) in reporter HeLa cells without and with 3mM histidinol 6 hours. WT cells with endogenous GCN2. GCN2 KO = knockout cells lacking functional GCN2. Variants expressed in the GCN2 KO lines are indicated. K619R = kinase dead negative control. n=3 biological replicates; 2-way ANOVA with group comparisons. (C) Representative immunoblots of GCN2 variant expression. Note – for clarity two blots are joined as indicated by vertical black lines between lanes 2&3, 3&4, and 11&12. (D) Quantification of C. n=4, 1-way ANOVA with multiple comparisons. * P<0.05, ** P<0.01, *** P<0.001, **** P<0.0001.

Sixteen GCN2 exemplar variants were selected as representative of each class of functional prediction, across a range of disease severities and distributed throughout the GCN2 protein (highlighted in Figure 1). Each variant was expressed in the GCN2-deleted reporter cells and bioluminescence was measured without or with histidinol treatment (Figure 2B). We noted a striking correlation between ATF4::NanoLuc reporter activation and diagnosis (Table 2). In all but one case, when reporter activation was preserved (P15L, I839T, T943A, L1148S, H1202Y), the clinical diagnosis had been of classical PAH, rather than either PVOD or PCH. The exception was Y34C, which had preserved reporter activation despite having been identified in an individual with PVOD. All other variants identified in PVOD or PCH showed impaired ATF4::NanoLuc responsiveness to histidinol.

**Table 2.**
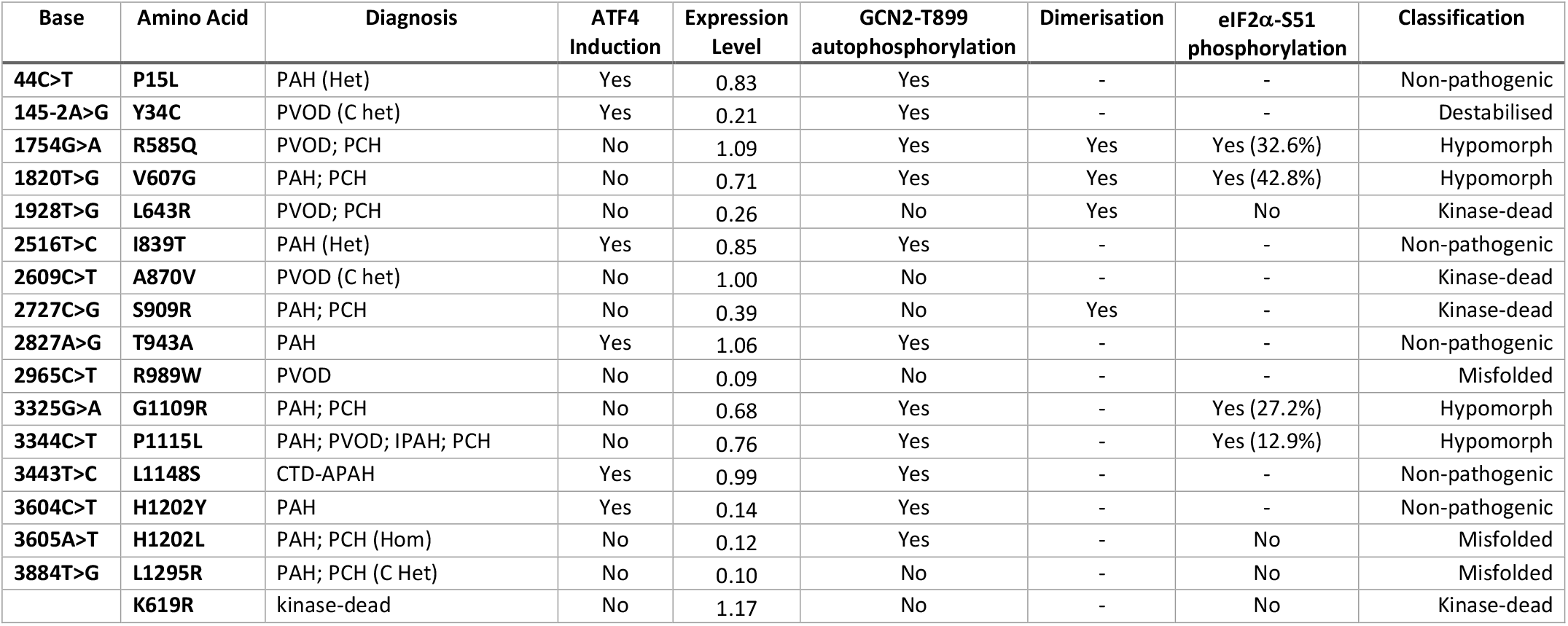
Variants analysis summary. ATF4 induction and protein expression from Figure 2. T899 autophosphorylation from Figure 3. Dimerisation from Figure 4. *In vitro* eIF2α phosphorylation from Figure 5.

| Base | Amino Acid | Diagnosis | ATF4 Induction | Expression Level | GCN2-T899 autophosphorylation | Dimerisation | eIF2α-S51 phosphorylation | Classification |
| --- | --- | --- | --- | --- | --- | --- | --- | --- |
| 44C>T | P15L | PAH (Het) | Yes | 0.83 | Yes | - | - | Non-pathogenic |
| 145-2A>G | Y34C | PVOD (C het) | Yes | 0.21 | Yes | - | - | Destabilised |
| 1754G>A | R585Q | PVOD; PCH | No | 1.09 | Yes | Yes | Yes (32.6%) | Hypomorph |
| 1820T>G | V607G | PAH; PCH | No | 0.71 | Yes | Yes | Yes (42.8%) | Hypomorph |
| 1928T>G | L643R | PVOD; PCH | No | 0.26 | No | Yes | No | Kinase-dead |
| 2516T>C | I839T | PAH (Het) | Yes | 0.85 | Yes | - | - | Non-pathogenic |
| 2609C>T | A870V | PVOD (C het) | No | 1.00 | No | - | - | Kinase-dead |
| 2727C>G | S909R | PAH; PCH | No | 0.39 | No | Yes | - | Kinase-dead |
| 2827A>G | T943A | PAH | Yes | 1.06 | Yes | - | - | Non-pathogenic |
| 2965C>T | R989W | PVOD | No | 0.09 | No | - | - | Misfolded |
| 3325G>A | G1109R | PAH; PCH | No | 0.68 | Yes | - | Yes (27.2%) | Hypomorph |
| 3344C>T | P1115L | PAH; PVOD; IPAH; PCH | No | 0.76 | Yes | - | Yes (12.9%) | Hypomorph |
| 3443T>C | L1148S | CTD-APAH | Yes | 0.99 | Yes | - | - | Non-pathogenic |
| 3604C>T | H1202Y | PAH | Yes | 0.14 | Yes | - | - | Non-pathogenic |
| 3605A>T | H1202L | PAH; PCH (Hom) | No | 0.12 | Yes | - | No | Misfolded |
| 3884T>G | L1295R | PAH; PCH (C Het) | No | 0.10 | No | - | No | Misfolded |
|  | K619R | kinase-dead | No | 1.17 | No | - | No | Kinase-dead |

Next, protein expression of the sixteen GCN2 variants was determined by immunoblotting (Figure 2C-D). Significantly reduced expression relative to wildtype GCN2 was observed for eight variants (Y34C, V607G, L643R, S909R, R989W, H1202Y, H1202L, L1295R). Of these, despite their low expression, two retained significant ATF4::NanoLuc reporter activation (Y34C and H1202Y). Conversely, three of these naturally occurring variants lacked reporter activity despite preserved expression (R585Q, G1109R and P1115L), similar to that seen for the artificial kinase-dead control (K619R). Of note, V607G showed no reporter activation despite only modestly reduced expression. Levels of GCN2 variants therefore do not fully reflect activity.

### GCN2 autophosphorylation is necessary but not sufficient for ISR induction

Autophosphorylation of GCN2 at T899 correlates with kinase activation in most circumstances^32,35^. Treatment with histidinol or starvation of amino acids, a physiological stimulus of the kinase, increased T899 phosphorylation of wildtype but not kinase-dead K619R GCN2 expressed in knockout cells (Figure 3A-B). When the naturally occurring variants were tested, eleven were capable of T899 autophosphorylation, while five were not (L643R, A870V, S909R, R989W, L1295R) (Figure 3C-F). Stressful stimuli triggered T899 autophosphorylation in all GCN2 variants capable of activating the ATF4::NanoLuc reporter, but also in five ISR-deficient variants (R585Q, V607G, G1109R, P1115L, H1202L) albeit only weakly (Figures 3C-F, summarised in Table 2).

**Figure 3.**
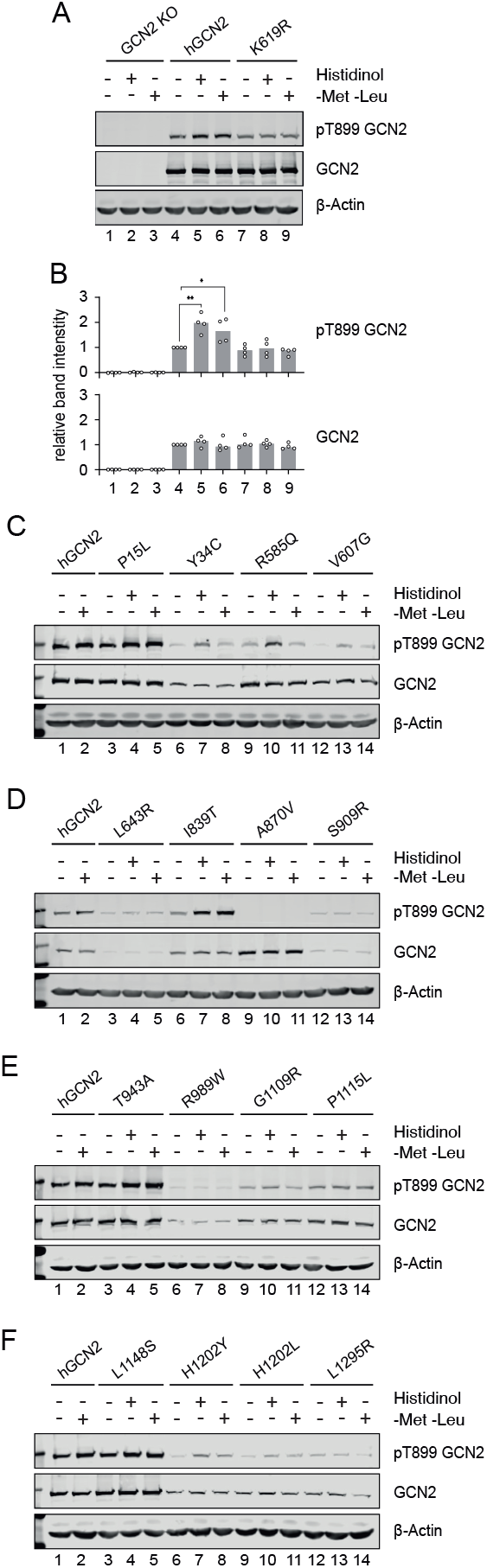
Stress-dependent activation of GCN2. (A) Representative immunoblot of GCN2 KO HeLa cells transiently transfected with constructs encoding full-length human GCN2 (hGCN2) or kinase-dead negative control (K619R). Hygromycin-selected pools were treated with either 7mM histidinol or starved of methionine and leucine (-Met -Leu) for 7 hours. (B). Quantifications of A. n=4 biological replicates normalised to untreated, hGCN2-transfected (lane 4). 1-way ANOVA with multiple comparisons (meaningful comparisons are shown). * P<0.05, ** P<0.01. (C-F). Representative immunoblots in GCN2 KO HeLa cells transiently transfected with GCN2 variants treat as in A. n=3 biological replicates.

Inactive GCN2 exists as an antiparallel homodimer and transitions to a parallel conformation on activation^36,37^. In yeast, stabilisation of the active state depends on the establishment of an intramolecular salt-bridge between residues R594 and D598, corresponding to R585 and E590 in the human protein^38^. The patient-derived R585Q variant was unable to activate the ATF4::NanoLuc reporter despite preserved expression (Figure 2) and autophosphorylation (Figure 3). Since the R585Q substitution is predicted to reposition the dimerisation interface salt-bridge, we sought to test if dimerisation was impaired. GCN2 constructs were generated tagged at the C-terminus with either 3xFlag or V5. Tagging did not impair GCN2 activity (Figure 4A). When co-expressed in GCN2 deleted cells, wildtype GCN2-3xFlag and GCN2-V5 formed mixed dimers detectable by anti-FLAG co-immunoprecipitation (Figure 4B). The R585Q variant similarly formed mixed dimers that could be co-immunoprecipitated (Figure 4C). The ISR-deficient kinase domain variants V607G, L643R, and S909R, predicted to have impaired folding (Table 1), were similarly able to form mixed dimers, suggesting that, at least for these variants, loss of dimerisation does not contribute to their impaired function (Figure 4C-F).

**Figure 4.**
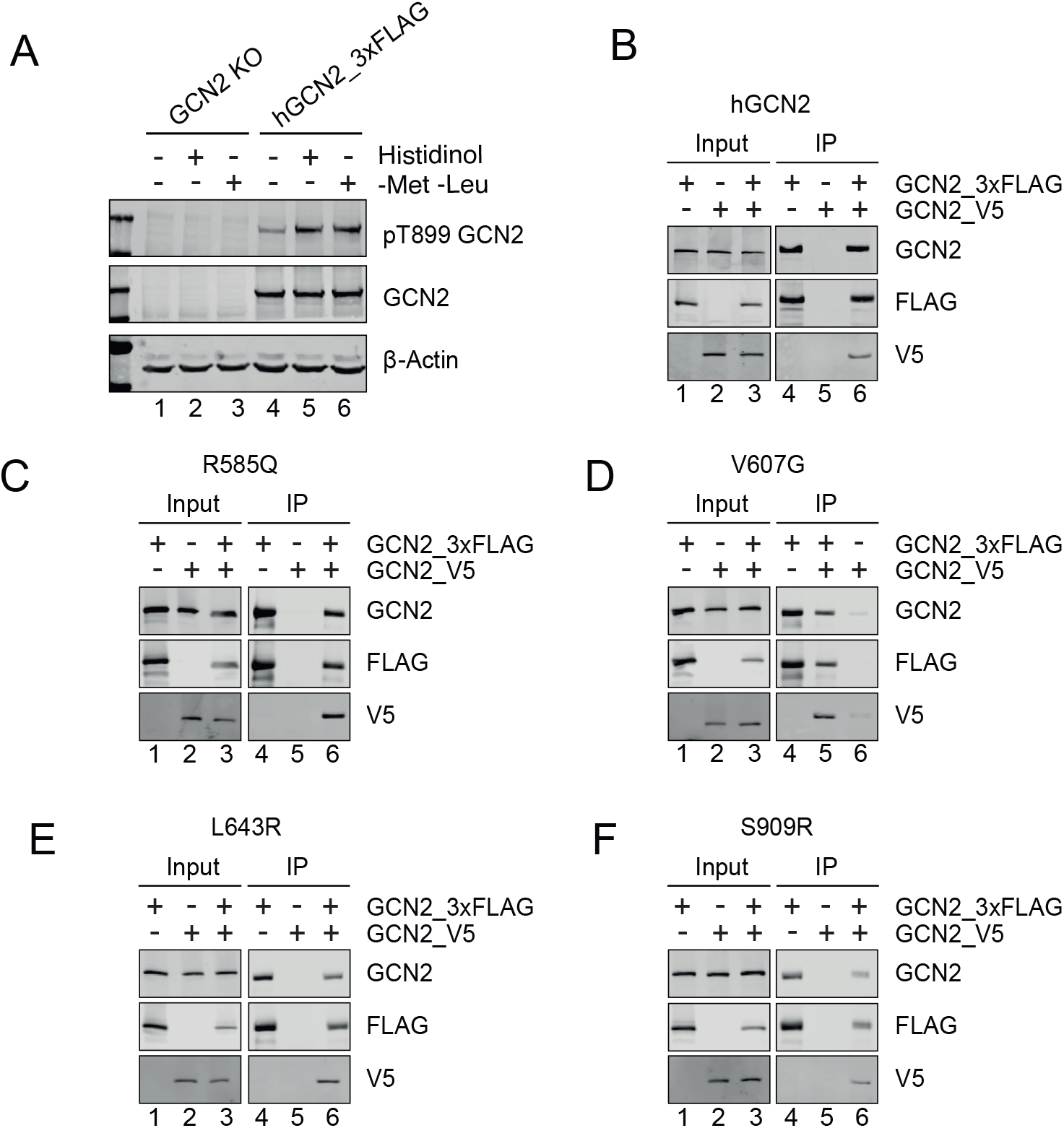
Dimerisation of variants of GCN2 variants. (A) Representative immunoblot of GCN2 KO HeLa cells transiently transfected with construct encoding 3xFlag tagged human GCN2 (hGCN2_3xFLAG) treated with either 7mM histidinol or starved of methionine and leucine (-Met -Leu) for 7 hours. n=3 biological replicates (B-F) Anti-FLAG immunoprecipitation-immunoblots from GCN2 KO cells transiently transfected with constructs encoding GCN2 variants tagged at the C-terminus with either 3xFLAG (GCN2_3xFLAG) or V5 (GCN2_V5). Lysates (lanes 1-3) and immunoprecipitates (lanes 4-6). Note co-immunoprecipitation of each pair of constructs consistent with dimer formation.

### Hypomorphic GCN2 variants can be activated by an ATP-competitive inhibitor

We then sought to test whether autophosphorylation-competent, but ISR-deficient variants might have lost the ability to engage the substrate eIF2α. Tagged GCN2 was recovered by immunoprecipitation of 3xFlag from lysates of cells starved of amino acids (Figure 5A), then immunopurified kinase was tested for its ability to phosphorylate the N-terminal domain of eIF2α (eIF2α-NTD) *in vitro* (Figure 5B-D). Wildtype but not kinase-dead K619R GCN2 showed enhanced T899 autophosphorylation when incubated with Mg-ATP, leading to increased phospho-GCN2 immunoreactivity and slower migration on SDS-PAGE (Figure 5B-D). When incubated with Mg-ATP and eIF2α-NTD, wildtype but not kinase dead K619R GCN2 phosphorylated eIF2α-NTD on serine 51 (Figure 5C-D). The autophosphorylation-competent variants R585Q, V607G, G11109R and P1115L (Figures 3C&E and 5B) also autophosphorylated their activation loop at T899 when incubated with Mg-ATP *in vitro*, and phosphorylated eIF2α-NTD albeit only weakly (Figure 5C-E). The variants L643R, S909R, L1202L and L1295R, which showed either weak or no autophosphorylation combined with low expression (Figure 5A&C) failed to phosphorylate eIF2α-NTD (Figure 5C-D). These data suggest that while some PVOD/PCH-associated variants preserve target engagement *in vitro*, their weak kinase activity appears insufficient for downstream signalling in cells. We classified these variants as hypomorphs.

**Figure 5.**
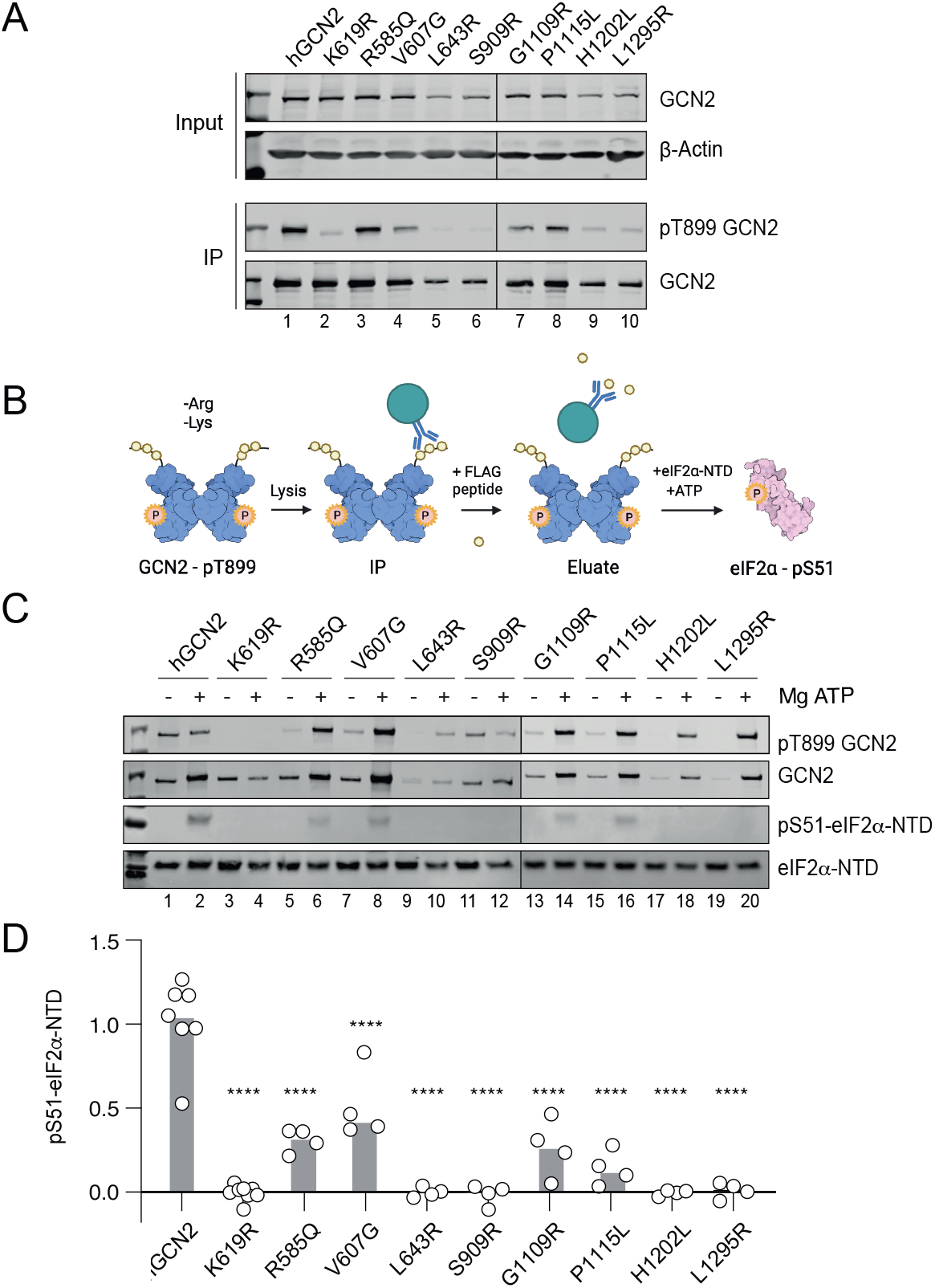
*In vitro* GCN2 kinase assay. (A) Representative Immunoprecipitation-immunoblots from cells expressing 3xFlag tagged GCN2. Cells were starved of arginine and lysine for 6 hours before harvesting. Intact GCN2 was eluted by competition via addition of excess FLAG peptide. Phosphorylation on threonine 899 reports autophosphorylation, note its absence for the K619R kinase-dead control. Note – for clarity two blots are joined as indicated by vertical black lines between 6&7. (B) Schematic of *in vitro* kinase assay drawn using Biorender.com. Tagged protein immunopurified using anti-FLAG beads then eluted with FLAG peptide. Bacterially-expressed recombinant eIF2α-N-terminal domain (NTD) served as a specific substrate. (C). Representative immunoblots of reaction products. Note – for clarity two blots are joined as indicated by vertical black lines between lanes 12&13. (D) Quantification of C; data presented with median value. n=4; 1-way ANOVA, compared to hGCN2-transfected control. **** P<0.0001.

It was recently shown how GCN2 could be activated paradoxically by sub-inhibitory concentrations of ATP-competitive kinase inhibitors^39,40^. Interestingly, the R585Q pathogenic variant, though not the L643R variant, was activated by treatment with the type-1.5 GCN2 kinase inhibitor, Gcn2iB^32^. We therefore tested the ability of such a small molecule, to activate the ATF4::NanoLuc reporter in cells expressing GCN2 variants. We found that each of the identified hypomorphs (R585Q, V607G, G11109R and P1115L) was rescued in their reporter activation by Gcn2iB, while the misfolded or kinase-dead variants were not (K619R, L643R and S909R; Figure 6).

**Figure 6.**
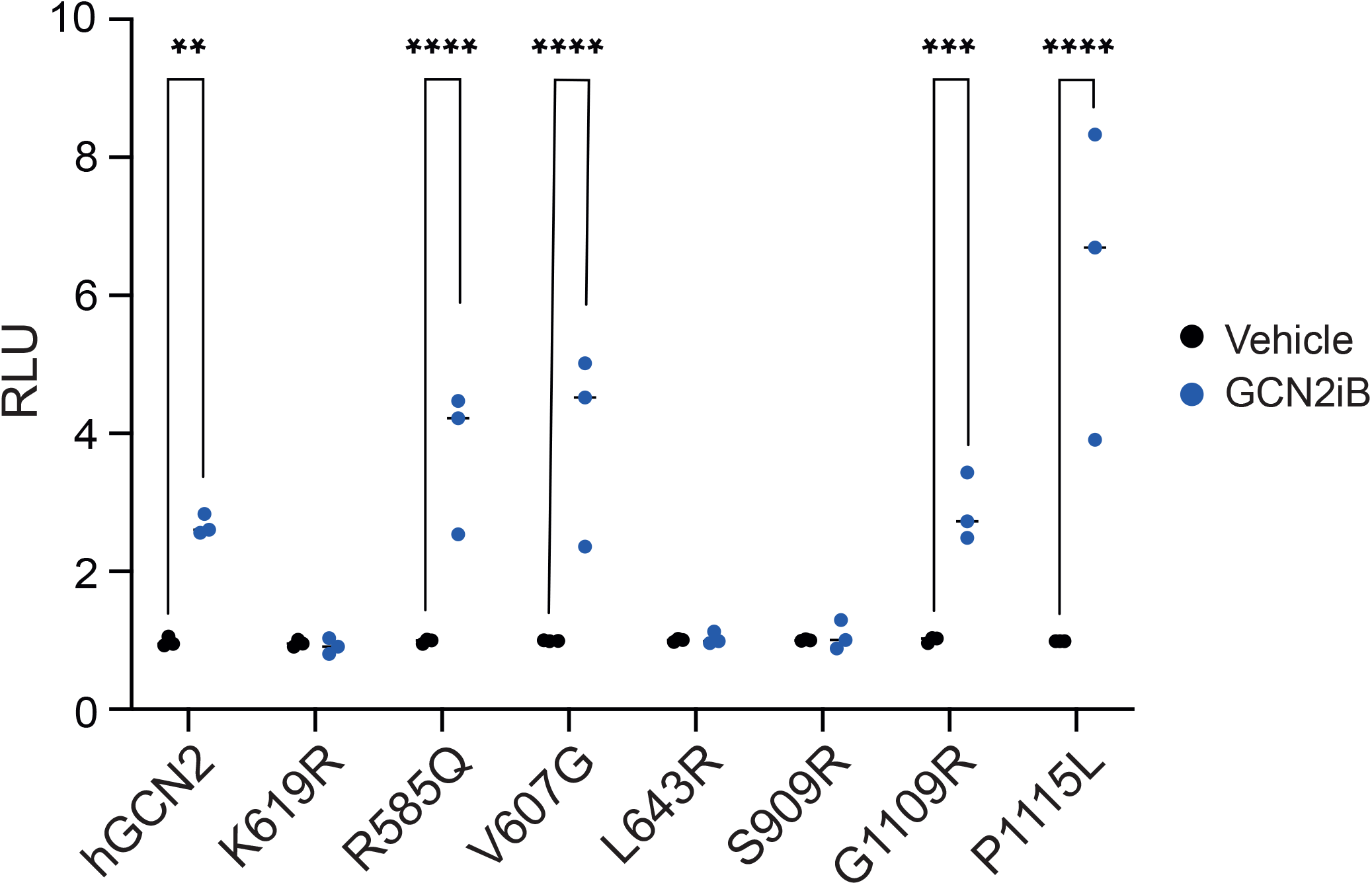
Activity of hypomorphic variants of GCN2 treated with Gcn2iB. ATF4::nanoLuc ISR reporter activity in GCN2 KO cells transiently transfected with constructions wild-type (hGCN2), kinase-dead (K619R), or selected patient variants. Cells were treated with 25nM Gcn2iB for 6 hours before luciferase assay. n=3 biological replicates; 2-way ANOVA with group comparisons. * P<0.05, ** P<0.01, *** P<0.001, **** P<0.0001.

## Discussion

Detection of biallelic pathogenic *EIF2AK4* mutations establishes the diagnosis of PVOD or PCH without the need for histological confirmation^6^. Validating the pathogenicity of *EIF2AK4* variants is therefore of significant diagnostic value. Segregation studies, although the gold standard, are not always feasible, in contrast to genetic sequencing, which is now routinely used in clinical practice. We found that computational analysis failed to identify some ISR-defective variants of GCN2. Our cell biological assay using a sensitive ATF4::nanoLuc reporter cell line is simple and inexpensive making it a feasible addition to characterisation workflows in specialist clinical practice.

We showcased this approach by evaluating sixteen of the thirty-four known missense variants of GCN2. Our observation that variants identified by genomic sequencing of individuals with classical PAH had preserved ISR reporter activity is consistent with GCN2 playing only a minor, or even no, role in that condition. Conversely, variants identified in individuals with either PVOD or PCH showed loss of ISR functionality, underlining the key role played by GCN2 in these disorders. The Y34C variant was a notable exception, maintaining some ISR reporter activity despite having been identified in an individual with PVOD patient. That patient also harboured a high-impact mutation of their second *EIF2AK4* allele (Lys190GlufsTer8, c.567dup). In our study, GCN2 Y34C was expressed at a significantly reduced level compared to the wild-type protein, raising the possibility that when combined with a null allele, the level of GCN2 generated might be insufficient to prevent development of the disease.

Importantly, we identified a subset of GCN2 variants with detectable but reduced kinase activity. These hypomorphs could be activated by the ATP-competitive inhibitor Gcn2iB, a type 11/2 inhibitor that stabilises the enzyme in a non-productive, yet active-like conformation (DFG-in, αC helix-out). It is believed that such binding to one protomer of a GCN2 dimer causes activation of the second drug-free promoter^39^. This suggests a potential therapeutic strategy for individuals with such hypomorphic variants identified by ATF4::nanoLuc reporter cells.

The experimental validation of GCN2 variants enabled us to evaluate computational predictive methods. It is recognised that evolutionary-based methods relying on homologous sequence alignment (e.g. SIFT, MAPP, PANTHER) are outperformed as stand-alone tools by approaches that integrate additional information such as structural features^41^. We compared the integrative tools PolyPhen2 and CADD, and found in our sixteen variants they incorrectly predicted enzyme activity in 5 and 6 instances respectively, giving a positive predictive value of approximately only 60%. Indeed, functional predictions using these methods are not recommended for diagnostic purposes for this reason^42^. Accuracy was improved with machine learning-based approaches, but even these rarely exceed 80% accuracy^43^. It has been estimated that 75% of disease-causing variants are linked to protein destabilisation^44^. Structure-based methods, such as the computation of Gibbs free energy variations, have also been attempted with FoldX recently being reported as the best performing method for the identification of disease-causing mutations^45^. FoldX predicted protein expression levels of GCN2 missense variants in our cell system, but the existence of stable but kinase-dead or hypomorphic variants limits its value. Nevertheless, integrating conservation-based approaches with structural features improves predictive performance. Meta-predictors, such as the ensemble method REVEL^28^, are better correlated with benchmark clinical datasets like ClinVar (reviewed in^46^). Although better than other tools, REVEL only returned 75% accuracy in our study. Recently AlphaMissense was developed, a deep-learning AlphaFold-derived system that combines residue structural context with unsupervised modelling of evolutionary constraints by comparing related sequences, claiming 90% accuracy in predicting the pathogenicity of missense variants when tested against the ClinVar dataset^31^. When using ISR activation as our gold standard, AlphaMissense incorrectly assigned 3 of the 16 GCN2 variants examined. The complexity of modelling protein-protein and protein-ribosome interactions adds to the challenge of using computational approaches to classify GCN2 variants^16,35^.

In summary, *in cellulo* evaluation of ISR activation using an ATF4::Nanoluc reporter outperformed existing computational approaches. This approach can identifies hypomorphic variants that can be revitalised by an ATP-pocket-binding small molecule drug.

## Materials and Methods

### Cloning and plasmids

All cloning and mutagenesis were performed via Gibson Assembly. A human codon-optimised GCN2 ORF was cloned into pcDNA3.1-Hygro(+) vectors and tagged with either 3xFLAG or V5 at the C-terminus. Site-directed mutagenesis was performed with specific primers for 16 naturally occurring variants and a kinase-dead control. A human ATF4 5’UTR ORF was inserted into pGL4.2 vectors and cloned in frame with Nluc-PEST® luciferase. A stop codon was inserted before the C-terminal degron to allow accumulation of the reporter.

### CRISPR/Cas9 knockout of *EIF2AK4* in HeLa cells

Human *EIF2AK4* specific guide RNAs were selected from the Brunello sgRNA library^47^. After primer duplex formation, guides were inserted in pSpCas9(BB)-2A-mCherry plasmids. Parental HeLa cells, cultured in DMEM supplemented with 10% FBS were transfected in 6-well dishes using 1μg DNA and Lipofectamine 2000 (1:3 ratio) in OptiMEM for 24 hours. At day 3 post-transfection mCherry-positive cells were sorted on a DB Melody cell sorter, as single cells into 96 well plates. Clones were screened by GCN2-targeting western blotting. Genetic mutations in KO clones were confirmed by genomic DNA extraction (100mM Tris, 5mM EDTA, 200mM NaCl, 0.25% SDS, 0.2mg/ml Proteinase K: incubation at 50°C overnight, then for 20 minutes at 98°C and clarification in a benchtop centrifuge at 10,000g for 5 minutes), PCR to amplify the locus targeted by the guide and subsequent NGS. Data were analysed using MacVector.

### Transfection and cell treatments

2x10^5^ HeLa cells were plated in 6-well dishes and let attach overnight before transfection. 1μg of plasmid DNA was mixed with Lipofectamine 2000 (1:3 ratio) in OptiMEM and incubated for 20 minutes at room temperature. 500μl of transfection medium were added onto the washed cells and topped up with additional 500μl of 10% FBS-DMEM. Transfection medium was removed after 24 hours. On day 2 cells were split and transfected cells were selected by 300μg/ml hygromycin treatment for 3 days. Cells were then maintained in 150μg/ml hygromycin 10% FBS-DMEM. Cell treatments were carried out as follows: 7mM histidinol for 7 hours (for western blotting); amino acid starvation in SILAC medium supplemented with 10% dialysed FBS and 25mM D-glucose (after 1x wash in PBS) without leucine only, leucine and methionine or lysine and arginine, for 7 hours.

### Immunoblotting

Cells were washed with PBS on ice and lysed in low-salt buffer (Buffer H: 10mM HEPES pH 7.9, 50mM NaCl, 0.5M sucrose, 0.1mM EDTA, 0.5% TX-100, 1mM DTT, 10ul/ml cOmplete™ Protease Inhibitor Cocktail (Roche, SKU 11836170001), phosphatase inhibitor cocktail mix (10 mM tetrasodium pyrophosphate, 17.5mM beta glycerophosphate, 100mM sodium fluoride), 1mM PMSF) and 10mM DTT. Samples were clarified at 4°C in a benchtop centrifuge at 10’000g for 10 minutes. Protein concentrations were estimated using Pierce BCA assay following the manufacturer’s instructions and equalised in lysis buffer and 6X loading buffer (1X: 60mM Tris-HCl pH 6.8, 10% glycerol, 2% SDS, 0.02% bromophenol blue, 1mM DTT). 8-14% polyacrylamide, 0.4% SDS gels were casted for SDS-PAGE. 60-80μg of protein were loaded per lane. Proteins were transferred to 25μm nitrocellulose membranes using 5% methanol transfer buffer. Blocking was done in 5% BSA in TBS as well as incubations with antibodies. Antibodies: GCN2 (in house, Ron lab: NY168); GCN2-phosphorT899 (Abcam, ab75836); β-actin (Abcam, ab8226); FLAG (Invitrogen, 740001); V5 (Abcam, ab27671).

### NanoGlo® luciferase assay

GCN2 KO, ATF4::NanoLuc reporter cells were transfected with GCN2 variant constructs as described above. Cells were seeded in 384-well plates at a density of 2.5x10^3^ cells per well in 20μl. Left-over cells were pelleted and lysed to confirm transfection efficiency via western blot. Wells were previously filled with 5μl of 15mM histidinol (5X) in 10% FBS-DMEM (final concentration 3mM), or 125nM (5X) Gcn2iB (final concentration 25nM). After 6 hours of incubation at 37°C and 5% CO_2_, 25μl of NanoGlo® luciferase assay reagent (buffer + substrate, 50:1) were added to the wells. Plates were mixed with orbital shaking for 1 minute and after 10 minutes of incubation at room temperature, bioluminescence signal (360-545nm) was acquired on a Tecan Spark plate reader.

### Dimerisation assay

GCN2 KO HeLa cells were transfected with 3xFLAG and/or V5 tagged GCN2 variant constructs as described above. After selection, cells were grown to confluence in 6cm dishes. Dishes were transferred on ice, washed once with PBS and lysed in 50mM Tris, pH 7.4, 150mM NaCl, 1mM EDTA, 1% Triton-X, plus protease and phosphatase inhibitors. Samples were clarified by centrifugation (>10,000g at 4°C for 10 minutes) and transferred in clean tubes on ice. Protein concentrations were measured using Bradford assay and equalised with lysis buffer. 80μg of protein samples were set aside as input controls, 600-800μg of protein were diluted in 200μl and added to 50μl of anti-FLAG® M2 affinity gel slurry (Millipore, A2220). Samples were incubated for 1 hour rotating at 4°C. Three washes were performed via centrifugation (1000rpm) using 1ml of lysis buffer without Triton-X. Elution was performed in 30μl of wash buffer plus 5μl of 1mg/ml 3xFLAG peptide solution (Pierce™ 3x DYKDDDDK, Thermo Scientific, A36805). Samples were incubated on ice for 15 minutes. Beads were pelleted by centrifugation (1000rpm). 15μl of eluate were used for western blotting.

### *In vitro* kinase assay

GCN2 KO HeLa cells were transfected with 3xFLAG GCN2 variant constructs as described above. After selection, cells were grown to confluence in 6cm dishes. Cells were washed once with PBS and incubated with amino acid depleted (lysine and arginine) SILAC medium supplemented with 10% dialysed FBS and 25mM D-glucose, for 6 hours. After immunoprecipitation (as described above) 10μl of eluate per reaction were added to PCR strip tubes or plates that were previously coated with 100mg/ml BSA for 3 hours, then washed 3X with PBS and dried thoroughly. Kinase assay was performed by adding recombinant eIF2α-NTD, amino acids 2-187 (kindly donated by the Ron lab) to a final concentration on 1μM and 500μM ATP in reaction buffer (1X: 50mM HEPES, pH 7.4, 100mM potassium acetate, 5mM magnesium acetate, 250μg/ml BSA, 10mM magnesium chloride, 5mM DTT, 5mM β-glycerophosphate). Reactions were incubated at 32°C for 10 minutes, then immediately quenched with 5μl of 6X SDS sample buffer for western blotting, pre-warmed at 95°C.

## Acknowledgements

Funding: SJM was supported by the MRC (MCMB MR/V028669/1 and MR/R009120/1), EPSRC (EP/R03558X/1), Cambridge Biomedical Research Centre (BRC-1215-20014); British Lung Foundation (BLF), Asthma+Lung UK (ALUK), Royal Papworth Hospital, and the Victor Philip Dahdaleh Foundation. GE were supported by philanthropic funding from Rick Medlock and by awards from EPSRC (EP/R03558X/1), BHF (RE/18/1/34212), Evelyn Trust (Grant 22/03), and the Cambridge-Tsinghua collaborative programme on sustainability and emerging technologies. JZ Doctoral Training Programme in Medical Research (DTP-MR; Cambridge Trust). This research was supported by the CIMR Flow Cytometry Core Facility. We wish to thank Reiner Schulte and Gabriela Grondys-Kotarba for their advice and support in flow cytometry cell sorting. We thank David Ron and Heather Harding for valuable discussions, advice, and reagents. We thank Glenn Masson for sharing reagents.

